# A reference genome sequence resource for the sugar beet root rot pathogen *Aphanomyces cochlioides*

**DOI:** 10.1101/2021.08.11.456025

**Authors:** Jacob Botkin, Ashok K. Chanda, Frank N. Martin, Cory D. Hirsch

## Abstract

*Aphanomyces cochlioides,* the causal agent of damping-off and root rot of sugar beet (*Beta vulgaris* L.), is a soil-dwelling oomycete responsible for yield losses in all major sugar beet growing regions. Currently, genomic resources for *A. cochlioides* are limited. Here we report a *de novo* genome assembly using a combination of long-read MinION (Oxford Nanopore Technologies) and short-read Illumina sequence data for *A. cochlioides* isolate 103-1, from Breckenridge, MN. The assembled genome was 76.3 Mb, with a contig N50 of 2.6 Mb. The reference assembly was annotated and was composed of 32.1% repetitive elements and 20,274 gene models. This high-quality genome assembly of *A. cochlioides* will be a valuable resource for understanding genetic variation, virulence factors, and comparative genomics of this important sugar beet pathogen.

## Genome Announcement

Sugar beets (*Beta vulgaris* L.) are grown globally and central to the multi-billion dollar sugar industry (Draycott, 2006). In 2020, the United States harvested 1.14 million acres of sugar beet, with a yield of 33.6 million tons valued at 1.6 billion dollars (USDA NASS, 2021). Beyond sugar production, sugar beets are utilized as a biofuel feedstock for ethanol production and the pulp byproduct is used to produce non-starch livestock feed (Alexiades et al., 2018). Aphanomyces root rot (ARR) of sugar beet occurs in all major growing regions and is especially devastating in years with above average precipitation (Draycott, 2006; Windels, 2000). The causal agent of ARR, *Aphanomyces cochlioides*, is a soil-dwelling oomycete capable of infecting sugar and table beets, spinach, chard, cockscomb, and numerous weed species (Grünwald & Coyne, 2003; Wen et al., 2006; Windels, 2000). Within the Saprolegniales, the genus *Aphanomyces* is composed of saprophytes, and pathogens of aquatic animals and plants (Makkonen et al., 2016; Martín-Torrijos et al., 2017; Wicker et al.,2001). *Aphanomyces cochlioides* produces thick-walled oospores that germinate zoosporangia and release infective zoospores (Islam, 2010). ARR is intractable due to long survival periods of oospores, which can persist in soil for several years (Windels & Brantner, 2000; Harveson et al., 2007). *Aphanomyces cochlioides* causes both acute damping-off, where the seedling hypocotyl becomes necrotic and constricted, and chronic ARR, which occurs throughout the growing season resulting in stunting, chlorosis, and necrotic water-soaked lesions on the taproot (Windels, 2000). In addition, severely diseased roots undergo sucrose losses post-harvest, adding to further economic loss (Campbell et al., 2006).

Currently, there are genome assemblies available for four *Aphanomyces spp.,* including *A. astaci*, *A. euteiches*, *A. invadans*, and *A. stellatus*. These resources have facilitated research in phylogenetics, evolution, comparative genomics, virulence factors, and diagnostic assays (Gaulin et al., 2018; Lévesque et al., 2010; Minardi et al., 2018). To date, there is no reference genome for *A. cochlioides*. We generated a *de novo* genome assembly for *A. cochlioides* using both long- and short-read technologies. This is the first reported *Aphanomyces spp.* genome sequenced with Nanopore long-read data, which resulted in a more contiguous assembly than other *Aphanomyces spp.* genomes. This resource will enable investigations into virulence mechanisms, evolutionary relationships, genetic diversity, and the development of specific detection assays.

*Aphanomyces cochlioides* isolate 103-1 (AC 103-1) was isolated from a field in Breckenridge, MN by soil baiting in 2014 followed by a single zoospore transfer on selective media (Windels, 2000). The isolate was maintained in sterile deionized ultrafiltered water at room temperature through transfers every six months. In 2020, this isolate was subjected to zoospore and oospore production assays, and spore structure and size were consistent with *A. cochlioides*. AC 103-1 zoospores were pathogenic on susceptible sugar beet cultivar Crystal 093 RR, with a high ARR disease severity index of 93.4 (Beale et al., 2002). High molecular weight (HMW) genomic DNA was extracted from 5-day old mycelium grown in 0.125X potato dextrose broth using a Genomic-tips 20/G kit (Qiagen, Inc., Valencia, CA, USA). The extraction procedure was modified by the addition of 5 mg Glucanex (Sigma-Aldrich, St. Louis, MO, USA) and 5 mg cellulase to Buffer G2 (Qiagen, Inc., Valencia, CA, USA), and an additional RNase A (Thermo Fisher Scientific, Inc., Waltham, MA, USA) treatment (dx.doi.org/10.17504/protocols.io.br3km8kw). The obtained HMW DNA was run on Oxford Nanopore Technologies MinION FLO-MIN111 flow cells on a GridION MK1, generating 24.4 Gb of long reads. The HMW DNA was also used to construct an Illumina TruSeq Nano DNA library and sequenced using a MiSeq v2 to generate 2 x 250 bp reads and a total of 6.38 Gb of sequence. All library preparations and sequencing were conducted by the Research Technology Support Facility at Michigan State University. The genome size was estimated to be ~82 Mb using the Illumina reads with Jellyfish v2.1.3 (Marçais & Kingsford, 2011). BEDtools v2.27.1 (Quinlan & Hall, 2010) was used to calculate the genome-wide coverage of short reads, which was estimated to be 77x.

To ensure the use of only high quality long-reads in the assembly the long-reads were filtered using NanoFilt v2.7.1 (De Coster et al., 2018) to remove reads shorter than 2 kb and a Phred quality score less than 8. Filtering retained 2,400,570 long-reads with an estimated genome coverage of 232x. Long-reads ranged from 2,000 - 349,045 bp, with a mean length of 7,696 bp. The filtered long-reads were *de novo* assembled using Canu v2.1 (Koren et al., 2017) with the options: ‘genomeSize = 82m’ ‘correctedErrorRate = 0.1’ ‘minOverlapLength = 500’ ‘minReadLength = 800’ ‘corOutCoverage = 100’. The assembly was polished with Racon v1.4.19 (Vaser et al., 2017) and Medaka v1.2.1 (https://github.com/nanoporetech/medaka). The Illumina reads were filtered to ensure high quality, a minimum length, and to remove adapter sequences using Trimmomatic v0.33 (Bolger et al., 2014). After filtering, 23,420,252 Illumina reads of 100 to 250 bp were used to error correct the polished assembly with Pilon v1.22 (Walker et al., 2014) in two iterations, resulting in 9,878 and 303 corrections, respectively (Andrews, 2010; Bolger et al., 2014). The final assembly size was 76.3 Mb, which is 92.7% of the estimated genome size of 82.4 Mb. The assembly contained 97 contigs with a contig N50 length of 2.6 Mb, a contig L50 of 11, and the longest contig was 5.0 Mb (Table 1, Figure 1). Assembly completeness was assessed using BUSCO v2.0 (Waterhouse et al., 2018) with the Alveolate-Stamenopiles lineage dataset of 234 benchmarking universal single-copy orthologs. The genome contained 93.2% of the ortholog groups with no duplicates. In addition, the completeness of the assembly was also assessed by mapping the Illumina reads to the assembly with Bwa-mem v0.7.17, which resulted in 99.988% of the reads having at least one alignment. Repetitive elements in the assembly were annotated with RepeatModeler v1.0.11 (Smit et al., 2008) and RepeatMasker v4.05 (Smit et al., 2013), which resulted in 32.1% of the assembly being annotated as repetitive elements (Table 1).

**Table 1.** Genome statistics of the *A. cochlioides* isolate 103-1 assembly and annotation.

| Feature | Statistic |
| --- | --- |
| Genome size estimation (Mb) | 82.4 |
| Genome assembly size (Mb) | 76.3 |
| Number of contigs | 97 |
| Longest contig (Mb) | 5.0 |
| Contig N50 (Mb) | 2.6 |
| Contig L50 | 11 |
| GC content (%) | 47.4 |
| Gene models | 20,274 |
| Protein-coding gene models | 19,514 |
| Total interspersed repeat content (%) | 32.1 |
| Complete BUSCOs (%) | 93.2 |

**Figure 1.**
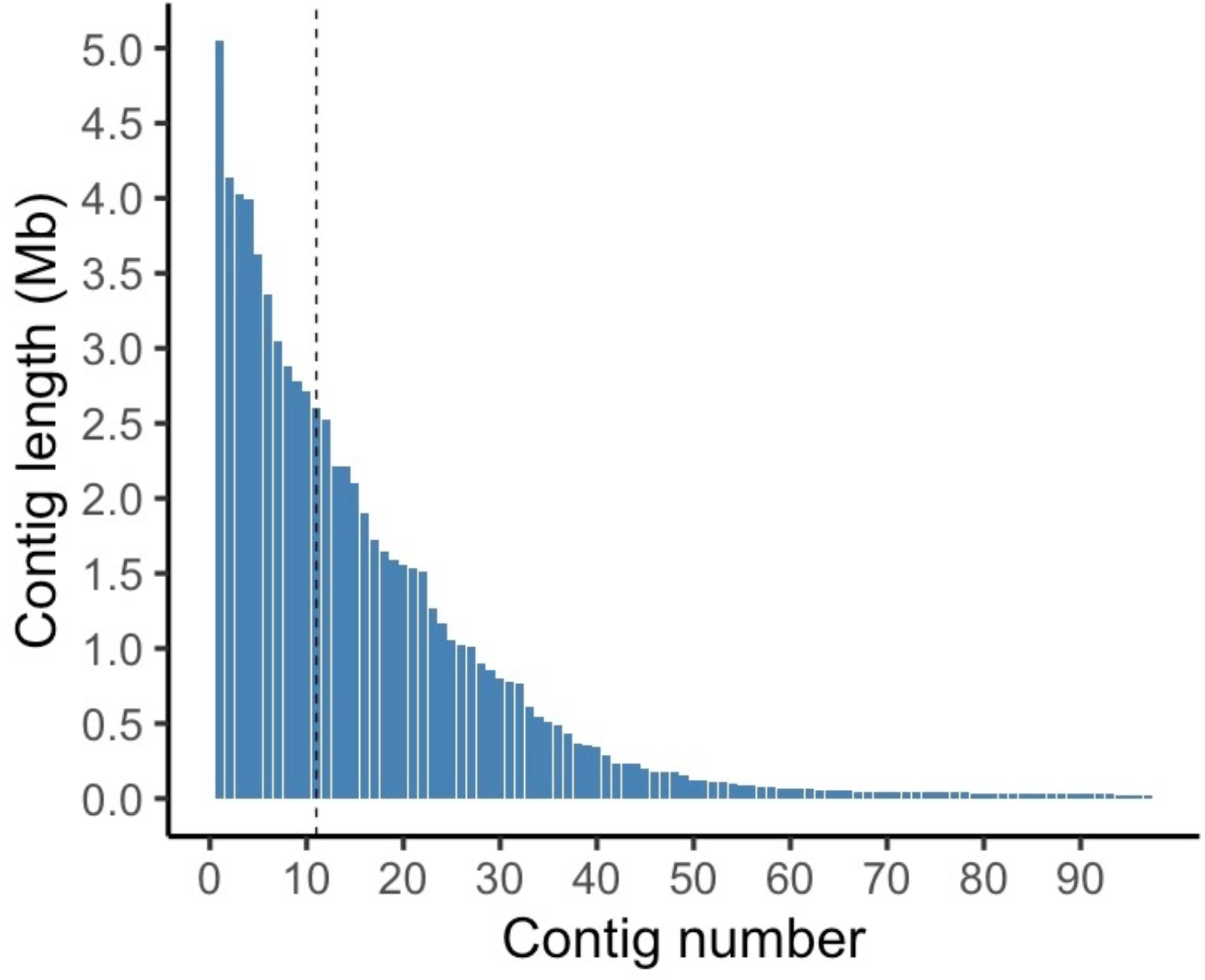
Length of contigs (Mb) in the final genome assembly ordered from longest to smallest. The black dashed line indicates the L50 contig.

To provide evidence for gene annotation RNA-seq data was generated from several different tissue types. Tissue from *A. cochlioides* oospores, mycelium, and infected sugar beet seedlings were collected, and RNA was extracted using a RNeasy Plant Mini Kit (Qiagen, Inc., Valencia, CA, USA). Mycelium was grown as described above, and oospores were grown on 2-week old sugar beet hypocotyls using established methods (Dyer & Windels, 2003). Also, 2-week old sugar beet seedlings of cultivar Crystal 093 RR were soil-drip inoculated with 5,000 zoospores per plant following established protocols, and diseased roots were removed from pots at 1, 3, 5, and 10 days post inoculation (dpi) for RNA extraction (Windels & Brantner, 2000). RNA from oospores, mycelium, and infected seedlings was sequenced using NovaSeq 2 × 150 bp sequencing, conducted by the University of Minnesota Genomics Center. RNA-seq reads were quality filtered with cutadapt v1.18 (Martin, 2011) at a minimum length of 40 bp and a minimum quality of 30, mapped to the *A. cochlioides* genome using STAR v2.5.3 (Dobin et al., 2013), and the mapped reads were used for genome annotation. The Funannotate v1.8.1 (Love et al., 2020) pipeline, consisting of funannotate scripts ‘clean’, ‘train’, ‘predict’, ‘update’, ‘annotate’, and ‘fix’ were used to predict and annotate gene models using the 172,894,133 mapped RNA-seq reads, 20,423 available *A. euteiches* protein sequences (Madoui et al., 2007), and 3,599 *A. cochlioides* mRNA sequences from NCBI. Within the assembled AC 103-1 genome, 20,274 gene models and 19,514 predicted protein-coding gene models were predicted. InterProScan v5.23 (Jones et al., 2014) was used to provide functional and GO annotation for predicted protein sequences.

All sequenced phytopathogenic oomycetes have been shown to contain CRN effectors, and some species, like *P. infestans*, contain RxLR effectors. A previous genomic analysis found that other pathogenic oomycetes in the Saprolegniales, like *A. euteiches, A. stellatus*, and *A. astaci* lack RxLR effectors (Gaulin et al., 2018). However, in our assembled AC 103-1 genome we identified 152 RxLR and 97 CRN effector candidates using the EffectR v1.02 package (Tabima & Grünwald, 2018) from predicted amino acid sequences generated by EMBOSS getorf (Rice et al., 2000). In addition, 1,594 secreted proteins were predicted by SignalP v4.1 (Nielsen et al., 2019). Of the predicted secreted proteins 702 were identified as effector candidates, including 280 apoplastic, 335 cytoplasmic, and 87 apoplastic/cytoplasmic effectors using EffectorP v3.0 (Sperschneider et al., 2018). This reference genome sequence could provide a resource for future research into virulence factors of *A. cochlioides*, comparative genomic studies, and phylogenomics.

## Resource availability

The genome assembly and annotation has been deposited in GenBank under the accession number JAHQRK000000000. All raw sequence data has been deposited in NCBI under BioProject PRJNA719397. BioSample SAMN18617838 contains Nanopore DNA sequences, Illumina DNA sequences, and the genome assembly and annotation. BioSamples SAMN18763738, SAMN18762525, SAMN18759881, SAMN18759656, SAMN18761261, SAMN18758017 contain RNA sequences. AC isolate 103-1 is available upon request to the corresponding authors and is stored at the University of Minnesota Bell Museum herbarium, St. Paul, MN. The generated and used in this project is available on GitHub at https://github.com/coryhirschlab/botkin_thesis/tree/main/chp2

